# Enhancing experience-dependent plasticity accelerates vision loss in a murine model of retinitis pigmentosa

**DOI:** 10.1101/2025.09.02.673825

**Authors:** Cecilia A. Attaway, Thomas C. Brown, Maureen A. McCall, Aaron W. McGee

**Affiliations:** Department of Anatomical Sciences and Neurobiology, School of Medicine; University of Louisville, Louisville, KY, 40202; Department of Ophthalmology and Visual Sciences, School of Medicine; University of Louisville, Louisville, KY, 40202; Department of Translational Neurosciences, School of Medicine, University of Arizona College of Medicine – Phoenix, Phoenix, AZ, 85004

## Abstract

Modulating neural plasticity is pursued as a therapeutic approach for several neurologic conditions. Here we evaluated if enhancing experience-dependent plasticity prolongs vision in a murine model of retinitis pigmentosa. First, we quantified the loss of visual acuity under both scotopic and photopic conditions for mice heterozygous for the P23H mutation in the *Rhodopsin* gene (*Rho P23H/+*). Acuity progressively declined under scotopic conditions followed by photopic conditions. Acuity deficits only broadly correlated with the retinal response measured by the electroretinogram. In contrast, acuity deficits were consistent with the percent of cortical excitatory layer 2/3 neurons responsive to higher spatial frequency visual stimuli. Then, we tested if enhancing plasticity in adult visual circuity by deleting the nogo-66 receptor gene (*Ngr1*) would preserve vision in *Rho P23H/+* mice. However, loss of vision was accelerated in *Ngr1 −/−; Rho P23H/+* mice. Thus, enhancing plasticity can be maladaptive in the context of neural degeneration.

## Introduction

Enhancing neural plasticity is pursued as a therapeutic intervention for neurodevelopmental disorders, neural injury, and neurodegenerative conditions (Cramer et al., 2011; LeBlanc and Fagiolini, 2011; Merzenich et al., 2014). In the visual system, sustaining or re-activating plasticity otherwise confined to a developmental critical period promotes recovery of acuity in rodent models of amblyopia (Hensch and Quinlan, 2018). Whether similar plasticity can prolong useful vision and acuity for other visual disorders including retinal degeneration is unclear. Testing this possibility may provide insight into the potential for modulating plasticity as a treatment for neurodegeneration.

Retinitis pigmentosa (RP) is a group of inherited retinal disorders that result in progressive loss of vision and has a worldwide prevalence of 1:4000 (Sohocki et al., 2001). RP preferentially disrupts the function of rod photoreceptor cells and RP patients are typically diagnosed in late adolescence to early adulthood following the onset of night blindness and impaired peripheral vision (Hartong et al., 2006). In the years following, vision continues to deteriorate and can result in low vision or complete vision loss (Ferrari et al., 2011). Longitudinal studies of patients with RP reveal that the loss of visual acuity is significantly slower than the deterioration of photoreceptors (Berson et al., 2002; Holopigian et al., 1996; Szlyk, 1997). This has led to the hypothesis that visual plasticity may play a role in preserving vision.

Approximately 30% of cases of RP are associated with an autosomal dominant (adRP) mutation (Daiger et al., 2015). More than 25 different genes and over 1000 mutations are associated with adRP, but the P23H point mutation in the *Rhodopsin* gene (*Rho*) is the most prevalent in North Americans, accounting for more than 10% of all cases (Daiger et al., 2015; Meng et al., 2020). The characteristic features of human P23H adRP are conserved in mice heterozygous for the *Rho P23H* mutation (Sakami et al., 2011). These mice exhibit a progressive decline in the full-field electroretinogram (ERG) that is associated with thinning of the outer nuclear layer where photoreceptors reside.

Visual acuity is constrained by the density of cones and retinal ganglion cells (RGCs), but also shaped by processing of visual information by visual cortex (Gianfranceschi et al., 1999; Prusky and Douglas, 2004). In the mouse, visual acuity can be measured with the visual water task (Prusky et al., 2000). This two-alternative forced choice test reveals that the typical visual acuity of wild-type (WT) adult mice under photopic conditions (photopic, ~100 cd/m2) is between 0.4 and 0.5 cycles per degree (cpd) (Prusky et al., 2000; Stephany et al., 2014). This acuity requires primary visual cortex (V1) (Prusky and Douglas, 2004). A fraction of excitatory neurons in V1 respond to sinusoidal gratings at spatial frequencies (SFs) slightly beyond behavioral estimates of visual acuity (~0.4-5 cpd) (Niell and Stryker, 2008; Salinas et al., 2017). Other studies have measured the opto-motor response, a reflex in rodents mediated by the superior colliculus, as a surrogate for visual acuity (Barwick et al., 2023). The spatial frequency thresholds measured for the optomotor reflex are lower than cortical-dependent visual tasks and mature faster than acuity measured with the visual water task (Prusky et al., 2004; Stephany et al., 2014). Overall, the relationship between impaired acuity, retinal function, and cortical function, has not been triangulated for any disorder of vision including RP.

The nogo-66 receptor 1 (NGR1) is one of several factors that limits plasticity in the adult brain (Stephany et al., 2016a). NGR1 is enriched at excitatory synapses (Lee et al., 2008; Loh et al., 2016). Adult ‘knock-out’ mice lacking a functional *Ngr1* gene (*Ngr1 −/−*) retain visual plasticity otherwise confined to the critical period and recover normal visual acuity in a mouse model of amblyopia, a prevalent developmental visual disorder (McGee et al., 2005; Stephany et al., 2018, 2016b, 2014). Here we established a framework for registering acuity to retinal function and cortical function for *Rho P23H/+* mice and then tested if enhancing visual plasticity by eliminating *Ngr1* expression would better sustain acuity during progressive retinal degeneration.

## Results

The visual water task measures functional vision. We modified this task to test and compare visual acuity under photopic and scotopic conditions. Photopic conditions were normal room light at 100 cd/m^2^. Scotopic conditions were 0.02 cd/m^2^. This scotopic condition, while not rod isolating, was the lowest condition under which the experimenter could manage this task. In addition, we tested acuity under monocular viewing conditions using the right eye. This eliminated the potential compensation by binocular vision, differences in acuity between the two eyes, as well as permitted direct comparisons of measurements of retinal and cortical function. Rods vastly outnumber cones in the mouse retina and represent nearly 97% of the photoreceptor population (Jeon et al., 1998). Thus, we anticipated that any deficits in visual acuity would be evident first under scotopic conditions where rod function predominates.

First, we measured photopic acuity (Fig. 1 and Suppl. Fig. 1). WT mice at ages ranging from 1-3 months (mo.) of age to 7-9 mo., displayed acuity in the typical range of 0.4 to 0.5 cpd (Prusky et al., 2000). WT mice at 10-12 mo. had slightly lower acuity (Fig. 1a). We then tested scotopic acuity and found that WT mice exhibited similar acuity under photopic and scotopic conditions at all ages (Fig. 1a).

**Figure 1.**
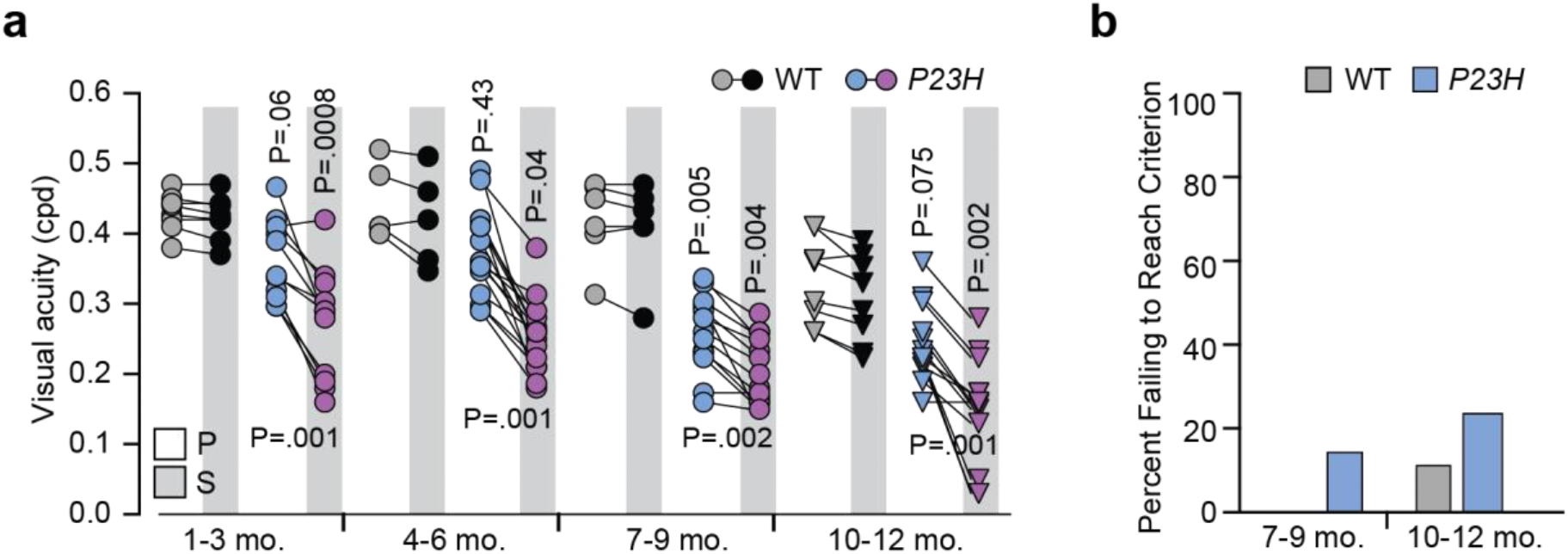
*Rho P23H/+* mice display a progressive deficit in acuity under both normal and low light conditions. (**a**) Visual acuity of WT and *Rho P23H/+* mice under photopic (P) and scotopic (S) conditions. Scotopic conditions are indicated by the grey bars. Lines connect the acuity measurements for each mouse under both conditions within an age group. P-values *above* symbols compare acuity of *Rho P23H/+* mice to age and luminance matched WTs (Brown-Forsythe and Welch ANOVA). P-values *below* pairs of symbols compare age groups of *Rho P23H/+* mice between photopic and scotopic conditions (mixed effects analysis). Tests corrected for all 16 comparisons between the two tests. (WT, P23H: 1-3 mo. n= 8,11; 4-6 mo. n=5,16; 7-9 mo. n=6,13; 10-12 mo. n=8,13). Circles and triangles represent individual mice (**b**) Fraction of mice under both photopic conditions that were unable to reach threshold criterion for testing of photopic visual acuity (WT, P23H: 7-9 mo. n= 0/6, 2/15; 10-12 mo. 1/9, 4/17).

By comparison, *Rho P23H/+* mice displayed significantly lower photopic acuity than age-matched WT mice at 7-9 mo.. The lower photopic acuity of WT mice at 10-12 mo. mitigated the difference between genotypes (Fig. 1a). *Rho P23H/+* mice also had significantly lower scotopic acuity than WT mice at all ages tested. As expected, the photopic visual acuity of *Rho P23H/+* mice was significantly higher than scotopic visual acuity across age. In addition to lower acuity, a fraction of 7-9 mo. and 10-12 mo. *Rho P23H/+* mice were unable to reach the criterion for testing, consistent performance at better than 70% at 0.05 cpd under photopic conditions, and therefore their visual acuity could not be assessed (Fig. 1b). In some cases, mice could not even orient towards the side of the tank displaying the visual stimulus. We set the acuity of these mice at 0.01 cpd for subsequent comparisons.

The prominent pre-clinical measurement of retinal dysfunction is the full-field ERG (Chang et al., 2002). The b-wave of the ERG represents the aggregate activity of ON bipolar cells that are post-synaptic to both rod and cone photoreceptors (Pinto et al., 2007). By 6 mo. of age, the photopic b-wave in *Rho P23H/+* mice is less than a third of the typical amplitude of WT mice and the scotopic b-wave is severely attenuated (Sakami et al., 2011). We measured the b-wave amplitude in response to full-field flashes for WT and *Rho P23H/+* mice at photopic and scotopic conditions that matched the parameters used to test visual acuity (Fig. 2). The *Rho P23H/+* mice at 7-9 mo. exhibited reductions in the photopic and scotopic b-waves (Barwick et al., 2023) (Fig. 2a-d). These deficits were exacerbated at 10-12 mo. and we could not record a photopic (4/15) or scotopic (11/15) b-wave in many *Rho P23H/+* mice (Fig. 2c,d).

**Figure 2.**
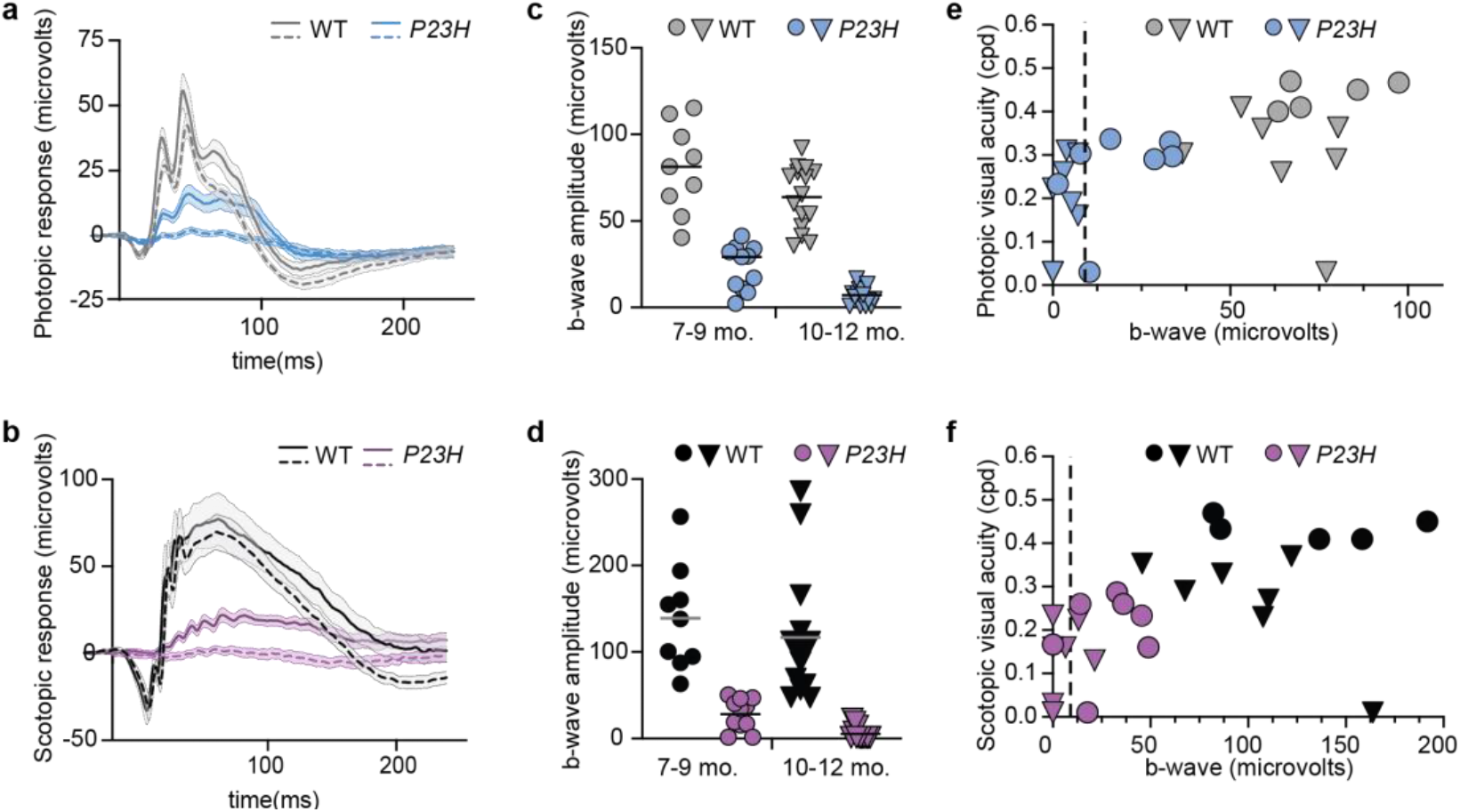
Reduced b-wave amplitude in *Rho* P23H/+ mice is associated with a range of acuities. **(a**,**b)** Averaged ± SEM ERG traces of WT (gray/black lines) and *P23H/+* mice (blue/purple lines) at 7-9 mo (solid lines) and 10-12 mo. (dashed lines) recorded for photopic flashes (16 dB) (panel a) and scotopic flashes (−20 dB) (WT, 7-9 mo. n=9, 10-12 mo. n=15; P23H, 7-9 mo. n=11, 10-12 mo. n=15) **(c**,**d)** Corresponding column plots of mean b-wave amplitude per mouse for photopic flashes (WT, gray; P23H, blue) and scotopic flashes (WT, black; P23H, purple). Circles represent individual mice at 7-9 mo and triangles mice at 10-12 mo. Horizonal lines indicate the mean. **(e**,**f)** Scatter plots of visual acuity and b-wave amplitude for the subset of WT and *P23H/+* mice with both measurements for photopic (panel e) and scotopic (panel f) conditions. Circles represent mice at 7-9 mo. and triangles at 10-12 mo. The vertical line represents mice with b-waves amplitudes less than 8 microvolts (WT, 7-9 mo. n=5, 10-12 mo. n=7; P23H, 7-9 mo. n=7, 10-12 mo. n=9).

Next, we compared photopic and scotopic acuity to the mean amplitude of their b-waves in a subset of mice for which both were measured (Fig. 2e,f). There was a clear separation between the distributions for WT and *Rho P23H/+* mice. Unexpectedly, the b-wave amplitude was a poor predictor of acuity. *Rho P23H/+* mice at 7-9 mo. and 10-12 mo. yielded photopic b-wave amplitudes that were ~10% or less of the average for WT mice yet possessed acuities ranging from unable to reach threshold for testing, up to.31 cpd, which overlapped with WT mice (Fig. 2e). A similarly broad range of acuities was associated with scotopic b-wave amplitudes (Fig. 2f).

Then, we compared photopic and scotopic visual acuity for WT and *RhoP23H/+* mice to the response properties of excitatory neurons in V1 evoked by visual stimuli presented under the same luminance conditions with 2-photon calcium imaging (Fig. 3 and Suppl. Fig. 2). We identified more than 3000 visually-responsive excitatory neurons in layer (L) 2/3 from WT and *Rho P23H/+* mice and calculated the orientation tuning and SF tuning for each. In WT mice, excitatory neurons in L2/3 span the full range of possible preferred orientations yet most prefer low SFs many octaves below acuity thresholds (Brown and McGee, 2025; Niell and Stryker, 2008; Salinas et al., 2017). Tuning for orientation was unremarkable as *Rho P23H/+* mice with impaired acuity exhibited distributions for preferred orientation and orientation selectivity index (OSI) similar to adult WT mice (Fig. 3a-b). In contrast, SF tuning was shifted to lower SFs in *Rho P23H/+* mice (Fig. 3c). In *Rho P23H/+* mice that did not reach criteria for testing on the visual water task, we were unable to identify populations of visually-responsive neurons (7 visually-responsive neurons from 434 segmented regions of interest (ROIs) and 3 mice).

**Figure 3.**
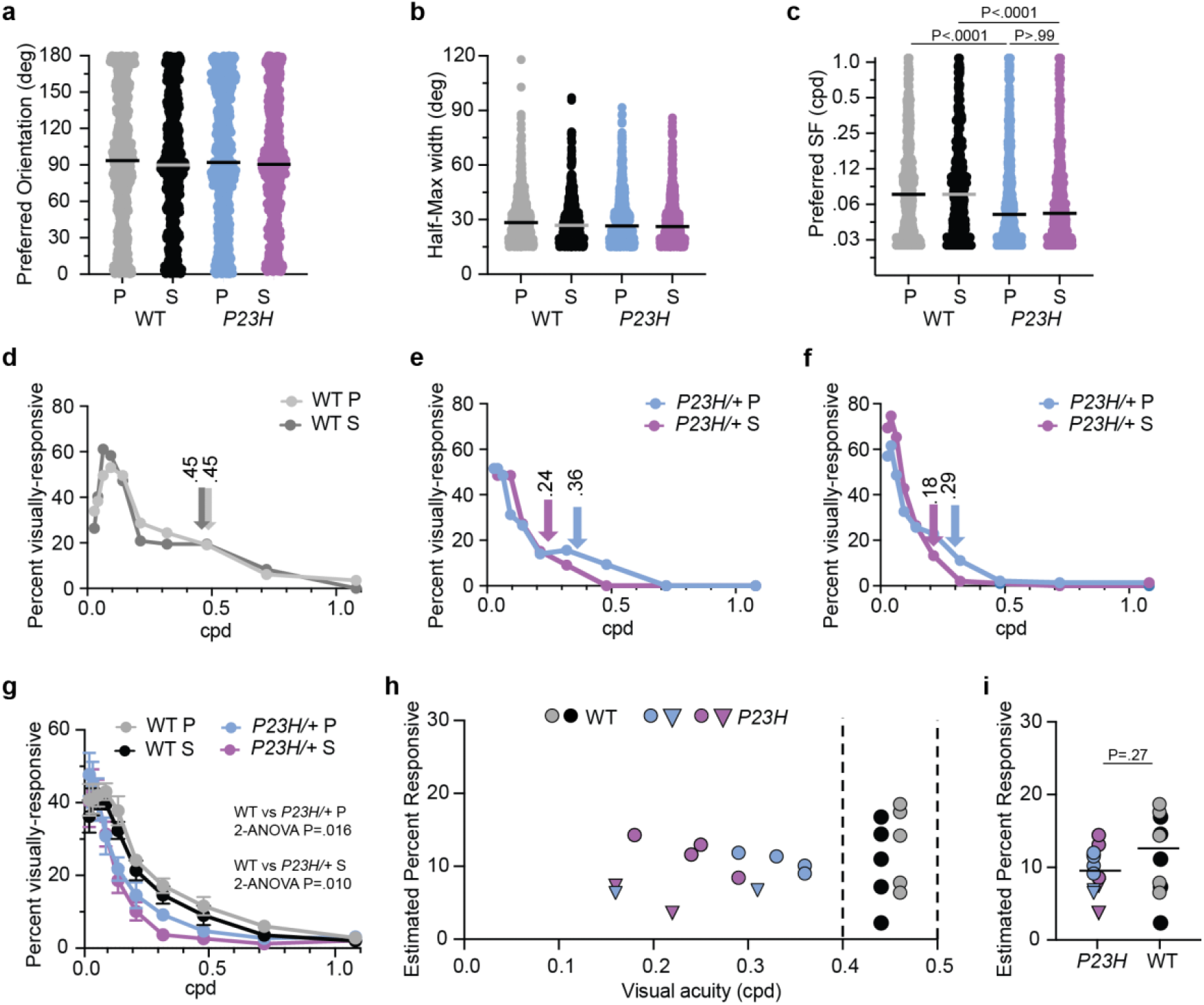
Cortical tuning to lower spatial frequencies is associated with impaired acuity in *Rho* P23H/+ mice. **(a)** Preferred orientation in degrees (deg), **(b)** half-maximum orientation tuning width (deg), and **(c)** preferred SF in cycles per degree (cpd) for excitatory neurons in L2/3 of V1 for WT (gray/black circles) and *P23H/+* mice (blue/purple circles) under photopic (P) and scotopic (S) conditions. Circles represent individual neurons. (WT, P n=800, S n=676; *P23H* P n=823, S n=729). Kruskal-Wallis test for (c) **(d-f)** Examples of SF response curves presenting the percent of visually-responsive neurons for each SF tested for a WT mouse **(d)** and two *rho P23H/+* mice **(e**,**f)** for both photopic (blue) and scotopic (purple) conditions. Arrows indicate the acuity measured for each mouse for the same luminance conditions as the calcium imaging. **(g)** SF response curves averaged across subjects for WT mice (n=5) and *P23H/+* mice (n=6) under photopic and scotopic conditions. Error bars represent standard error of the mean (SEM). Two-way ANOVA comparing WT and *P23H/+* mice at each luminance condition **(h)** Interpolated percent of visually-active neurons for each mouse in panel g responding at the SF corresponding to the measured visual acuity as in panels d-f. Values for WT mice are presented between dashed vertical lines bound the range of acuities for 1-3 mo. WT mice presented in Fig. 1a. Circles represent individual mice at 7-9 mo and triangles mice at 10-12 mo. for *P23H/+* mice. **(i)** column plot for the data presented in panel h. Unpaired two-tailed t-test.

At the population level, the preferred SF tuning distributions were similar for photopic and scotopic conditions for *Rho P23H/+* mice, although each *Rho P23H/+* mouse had lower acuity under scotopic conditions (Fig. 1a and Fig. 3c). This is likely the consequence of the considerable overlap of the range of acuities under the two luminance conditions for *Rho P23H/+* mice older than 7 mo. of age (Fig. 1a). Therefore, to more accurately determine how acuity related to the SF tuning of populations of V1 neurons, we plotted the fraction of visually-responsive neurons at each SF tested for individual WT and *Rho P23H/+* mice with their acuity for both photopic and scotopic conditions (Fig. 3d-h). Representative examples of WT and *Rho P23H/+* mice reveal that SF response curves are shifted towards lower SFs in *Rho P23H/+* mice with lower acuity (Fig. 3d-f). Averaging these SF response curves across mice revealed significant differences between genotypes (Fig. 3g). We also estimated by linear interpolation from these SF response curves for each mouse the percent of neurons responsive at the acuity measured behaviorally (Fig. 3 d-f,h,i). The scatter plot reveals that the estimated fraction of neurons responsive at the acuity threshold resides near 10 percent of the visually-responsive population for both adult naive WT mice and *Rho P23H/+* mice (Fig. 3h,i).

Last, given this relationship between acuity and visual cortical function, we tested the hypothesis that increased plasticity in visual circuitry would sustain acuity in this mouse model of adRP. We generated *Ngr1 −/−; Rho P23H/+* mice and measured their acuity followed by measurements of retinal function with the ERG, the tuning properties of neurons in V1 by calcium imaging, and thickness of the retinal outer nuclear layer by histology, at 7-9 mo. and 10-12 mo. of age (Fig. 4). The following results refuted this hypothesis.

**Figure 4.**
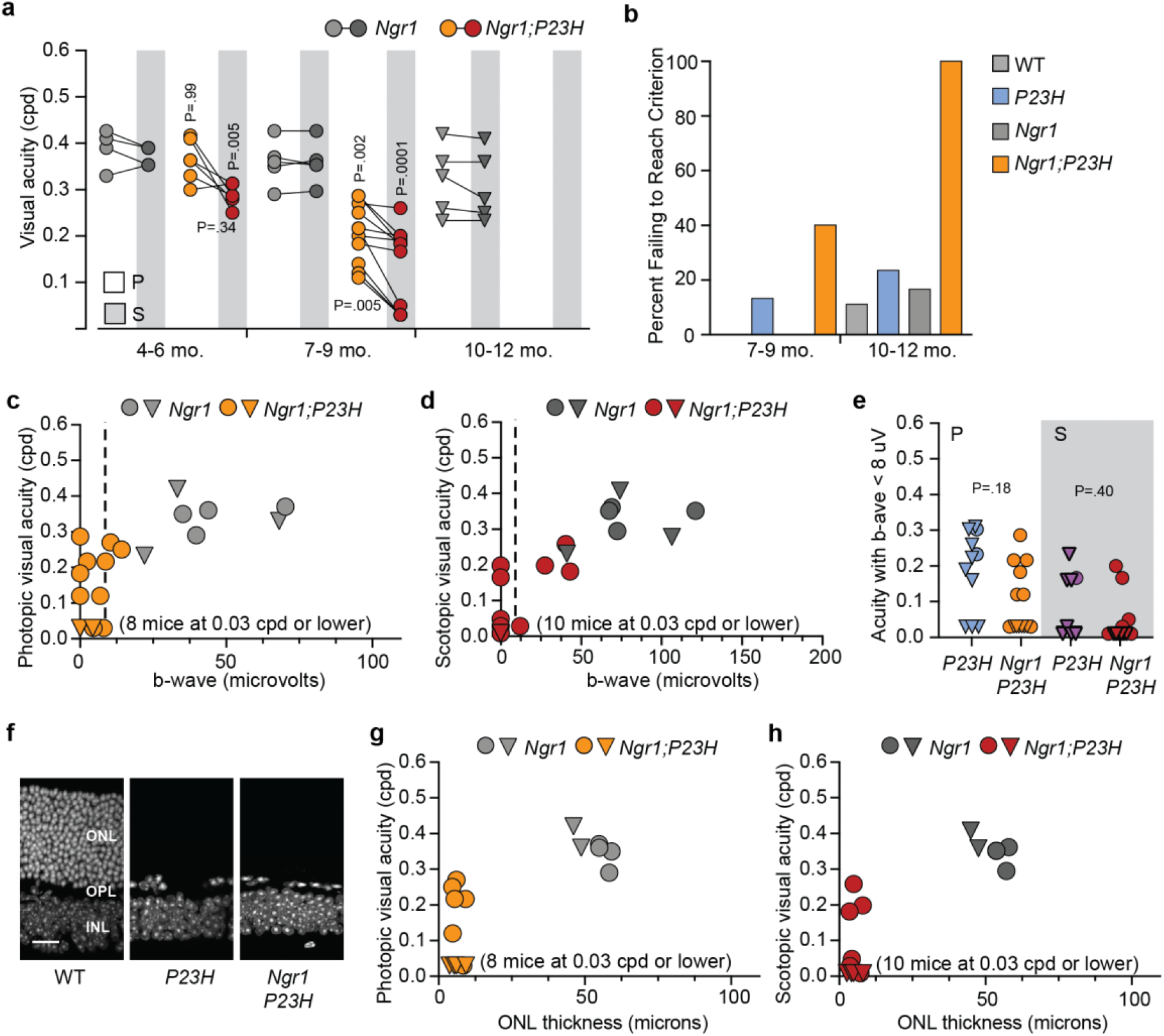
Enhancing visual plasticity accelerates loss of vision in *Rho P23H/+* mice. **(a)** The visual acuity of *Ngr1 −/−* and *Ngr1 −/−; Rho P23H/+* mice under photopic (P) and scotopic (S) conditions. Scotopic conditions are indicated by the grey bars. Lines connect the acuity measurements for each mouse under both conditions within each group. P-values above symbols compare *Ngr1 −/−; Rho P23H/+* mice to age and luminance matched *Ngr1 −/−* group (Brown-Forsythe and Welch ANOVA). P-values below symbols compare *Ngr1 −/−; Rho P23H/+* mice between P and S luminance conditions (mixed effects analysis). Tests corrected for all 6 comparisons between the two tests. *Ngr1 −/−* + *Rho P23H/+*: 4-6 mo. n=4,6; 7-9 mo. n=5,12; 10-12 mo. n=5,7 **(b)** Fraction of mice under photopic conditions that were unable to reach threshold criterion for testing. WT and *P23H/+* are reproduced from Figure 1a.: *Ngr1 −/−* + *Rho P23H/+*: 7-9 mo. n= 0/5, 8/20; 10-12 mo. 1/6, 7/7. **(c**,**d)** Scatter plots of visual acuity and b-wave amplitude for the subset of *Ngr1 −/−* and *Ngr1 −/−; Rho P23H/+* mice with both measurements for photopic (panel c) and scotopic (panel d) conditions. Circles represent mice at 7-9 mo. and triangles at 10-12 mo. The vertical line represents mice with b-waves amplitudes less than 8 microvolts (*Ngr1 −/−*, 7-9 mo. n=4, 10-12 mo. n=3; *Ngr1 −/−; Rho P23H/+*, 7-9 mo. n=10, 10-12 mo. n=6). **(e)** Column plot of acuity values for mice with b-wave amplitudes less than 8 microvolts. WT and P23H are reproduced from Figure 1 e,f. Kruskal-Wallis test. **(f)** Nissl-stained histological cross sections of retina from WT, *Rho P23H/+*, and *Ngr1 −/−; Rho P23H/+* mice at 10-12 mo.. The positions of the outer nuclear layer (ONL), Outer plexiform layer (OPL) and Inner nuclear layer (INL) are shown. Scale bar = 20 microns. (**g**,**h**). Scatter plots of visual acuity and ONL thickness for the subset of *Ngr1 −/−* and *Ngr1 −/−; Rho P23H/+* mice with both measurements for photopic (panel g) and scotopic (panel h) conditions. Circles represent mice at 7-9 mo. and triangles at 10-12 mo. (*Ngr1 −/−*, 7-9 mo. n=4, 10-12 mo. n=2; *Ngr1 −/−; Rho P23H/+*, 7-9 mo. n=6, 10-12 mo. n=4).

The maturation of acuity by *Ngr1 −/−* mice is normal (Stephany et al., 2014). Acuity of *Ngr1 −/−* mice at 10-12 mo. was similar to WT mice (Fig. 4a and Suppl. Fig. 3a-b). *Ngr1 −/−; Rho P23H/+* mice displayed a reduction in acuity under scotopic conditions similar to *Rho P23H/+* mice at 4-6 mo. of age (Fig. 4a). However, the deficits in acuity were significantly greater in *Ngr1 −/−; Rho P23H/+* mice than *Rho P23H/+* mice at 7-9 mo. and older (Fig. 4a and Suppl. Fig. 4a-b). In addition, the fraction of mice unable to reach the criteria for testing on the visual water task doubled for *Ngr1 −/−; Rho P23H/+* mice relative to *Rho P23H/+* mice at 7-9 mo. (Fig. 4b). Critically, none of the *Ngr1 −/−; Rho P23H/+* mice examined could reach criterion at 10-12 mo.. This differed dramatically from the performance of *Rho P23H/+* mice at this age. Thus, enhancing plasticity accelerated the loss of vision in this model of RP.

This loss of vision could not be attributed to a commensurate decline of retinal function in *Ngr1 −/−; Rho P23H/+* mice. The b-wave amplitude was only slightly less than age-matched *Rho P23H/+* mice under both photopic and scotopic conditions. Mice with b-wave amplitudes near the noise level spanned the same broad range of acuity between genotypes (Fig. 4 c-d and Suppl. Fig. 3c-d). Plotting the photopic and scotopic acuity for mice with b-wave amplitudes less than 10% of the control mean (~8 microvolts) did not support a significant difference between groups (Fig. 4e).

Adult *Ngr1 −/−* mice display tuning for orientation and spatial frequency indistinguishable from WT mice (Brown and McGee, 2025). We performed calcium imaging on *Ngr1 −/−; Rho P23H/+* mice that expressed GCaMP6s; however, these functionally blind mice did not yield enough visually-responsive neurons for analysis (46 visually-responsive neurons from 848 ROIs and 3 mice) (Suppl. Fig. 4).

The number and function of photoreceptors dictate the upper limit of vision in RP. The amplitude of the ERG b-wave reflects the cumulative signaling from photoreceptors to ON bipolar cells, and differences in the number of photoreceptors might be masked in the b-wave amplitude. Therefore, we measured and compared the thickness of the outer nuclear layer (ONL) in 7-12 mo. *Ngr1 −/−* and *Ngr1 −/−; Rho P23H/+* mice (Fig. 4 f-h). The *Ngr1 −/−; Rho P23H/+* ONL matched the range of thickness reported for *Rho P23H/+* mice at 6 mo. of age and older (Barwick et al., 2023; Lobanova et al., 2018; Sakami et al., 2011; Wang et al., 2023). Some mice with ONLs that had an average thickness corresponding to a single layer of photoreceptors (or less) possessed photopic and scotopic acuity better than 0.20 cpd, while others with equally thin ONLs were functionally blind (Fig. 4 g-h). Thus, we conclude that an increased loss of photoreceptors is not the primary cause of vision loss in *Ngr1 −/−; Rho P23H/+* mice.

## Discussion

In summary, we triangulated the progressive loss of visual acuity in the *Rho P23H/+* mouse model of RP with retinal and cortical function (Sakami et al., 2011). Deficits in acuity were greater under scotopic conditions across all ages for *Rho P23H/+* mice. The severity of acuity impairment correlated only broadly with deficits in retinal function measured with full-field ERG but were consistent with the percentage of neurons in primary visual cortex responsive to higher SFs measured with calcium imaging. Contrary to our hypothesis, enhancing neural plasticity was deleterious for visual function in *Rho P23H/+* mice.

The full-field ERG is an effective tool for identifying retinal dysfunction but lacks the sensitivity to discriminate the severity of visual deficit. The pattern ERG may provide a better estimate of the loss of responsiveness to higher SFs, but the magnitude of the response is only a fraction of the flash ERG (less than 8 microvolts at 0.1 cpd) and suffers from the same limitations in the threshold for detection, particularly for mice with photoreceptor degeneration (Porciatti, 2007). Cortical visually-evoked potentials (VEPs) have also been employed to estimate visual acuity (Domenici et al., 1991). However, VEPs represent a complex combination of synchronous aggregate subthreshold and spiking activity (Einevoll et al., 2013). In rodent models of amblyopia, acuity estimated by VEPs can be more than two octaves higher than acuity measured with the visual water task (He et al., 2007). By comparison, we calculated the percent of visually responsive neurons in L2/3 of V1 for *Rho P23H/+* mice with a range of acuity deficits and determined that ~10% of neurons responded at the SF corresponding to the behavioral acuity limit. This percent of visually responsive neurons may reflect the size of the cortical population required for perception. It would be interesting to ascertain if increasing the excitability of neurons tuned to higher SF might improve acuity as demonstrated for neural ensembles in head-fixed assays of contrast perception (Bounds and Adesnik, 2025).

How might enhancing plasticity in visual circuitry accelerate the loss of acuity in this model of retinal degeneration? Expression of *Ngr1* gene is not detectable in photoreceptors, but is present along the visual pathway in retinal ganglion cells (RGCs), visual thalamus, and visual cortex (Solomon et al., 2018; Stephany et al., 2018). Perhaps the progressive deterioration of photoreceptors provides a signal that attenuates the synaptic drive onto RGCs or the organization of the retinogeniculate inputs to visual thalamus. Expression of *Ngr1* in thalamus limits plasticity that permits recovery of acuity in a murine model of amblyopia upon the restoration of normal (binocular) vision after a month of monocular deprivation (Stephany et al., 2018). This plasticity may operate in reverse during progressive retinal dysfunction to decimate acuity. Alternatively, cortical plasticity may be the culprit. Recently, we discovered that adult *Ngr1 −/−* mice retain the high degree of representational drift in visual cortex that is otherwise confined to the developmental critical period (Brown and McGee, 2025). An increased capacity for adult excitatory cortical neurons to adapt their tuning properties to recent visual experience may permit cortical neurons to adopt non-visual inputs to more rapidly replace degrading feed-forward visual drive as their major source of excitation, thereby accelerating the loss of functional vision. Future work will be required to discriminate between these possibilities.

The visual system is a useful model for investigating the role of plasticity in neurodegeneration. Both behavioral performance and neuronal function can be readily evoked with quantitative visual experience. Retinal circuitry is well characterized, and many of the tools employed for measuring retinal function clinically are also available for rodent models. Neuronal function can be measured independently along several stages of the visual pathway. Last, vision is not required for viability so even severe dysfunction can be observed. We conclude that enhancing plasticity is deleterious for vision in the context of retinal degeneration. Based on these findings, we propose that experience-dependent plasticity is predominantly maladaptive for neural circuits during neurodegeneration.

## Methods

**Table 1.** Reagents and resources employed in the study.

| REAGENT or RESOURCE | SOURCE | IDENTIFIER |
| --- | --- | --- |
| **Deposited data** |  |  |
| Calculated tuning properties for all neurons |  | N/A |
| **Experimental models: Organisms/strains** |  |  |
| Mouse: B6;DBA-Tg(tedO-GCaMP6s)2Niell/j | The Jackson Laboratory | RRID: ISMR_ JAX: 024742 |
| Mouse: B6;Cg-Tg(Camk2a-tTA)1Mmay/DboJ | The Jackson Laboratory | RRID: ISMR_ JAX: 007004 |
| Mouse: B6.129S6(Cg)-Rhotm1.1Kpal/J | The Jackson Laboratory | RRID:IMSR_ JAX:017628 |
| Mouse: nogo-66 receptor (ngr1/rtn4r) constitutive mutant mice |  | Kim et al, 2004 |
| **Software and algorithms** |  |  |
| MATLAB | Mathworks | https://www.mathworks.com/ |
| Processing2 | Processing | https://processing.org/ |

### Lead contact

Further information and requests for resources and reagents should be directed to and will be fulfilled by the Lead Contact, Aaron W. McGee.

### Materials availability

This study did not generate new unique reagents

### Experimental model and subject details

All procedures were approved by University of Louisville Institutional Animal Care and Use Committee (IACUC) protocol 22105 and were in accord with guidelines set by the US National Institutes of Health. Mice were anesthetized by isoflurane inhalation and killed by carbon dioxide asphyxiation or cervical dislocation following deep anesthesia in accordance with approved protocols. Mice were housed in groups of 5 or fewer per cage in a 12/12 light–dark cycle. Animals were naive subjects with no prior history of participation in research studies. A total of 195 mice, both male (100) and female (95) were used in this study.

### Mice

*Rho P23H/+* mice (stock no. 017628) were purchased from Jackson labs and crossed to double transgenic mice, Tg-CaMKII-tTA (stock no. 007004) and Tg-TRE-GCaMP6s (stock no. 024742), to express GCaMP6s in forebrain excitatory neurons (Mayford et al., 1996; Wekselblatt et al., 2016) Mice were genotyped with primer sets suggested by Jackson Labs. These mice were crossed onto the *Ngr1* −/−background (Kim et al., 2004). Mice were genotyped with custom primer sets.

### Monocular Lid Suture

To measure visual acuity from the right eye, the left eyelid was sutured shut using a single mattress suture made of 6-0 polypropylene monofilament (Prolene 8709H; Ethicon) under brief isoflurane anesthesia (2%). The knot was secured with cyanoacrylate glue. The eyelid suture was removed when the visual water task was fully completed.

### Visual Water Task

The Visual Water Task is a two-alternative, forced-choice discrimination task that measures cortically mediated visual acuity (Prusky et al., 2000). In brief, a monitor was positioned at the wide end of a custom made trapezoidal shaped tank behind clear plexiglass. An opaque divider (46 cm long) divided the monitor and created a point where the mouse had to choose which side to swim towards. One side of the screen displays a sinusoidally modulated grating at 95% contrast. The other side of the screen displays an isoluminant gray screen. The spatial frequency of the grating was calculated from the screen to the choice point (46cm). The tank was filled with room temperature water, and a hidden platform was submerged below the water’s surface in front of the monitor displaying the grating.

Mice were divided into two groups (flights) (≤ 6 mice per flight, total of ≤ 12 mice in a cohort) with alternating rest and test blocks. Mice were trained to swim towards the monitor with the low frequency (0.05 cycles per degree) grating and the hidden platform. The side on which the stimulus was displayed was pseudorandom. In a successful trial, the mouse swam past the midline divider to the monitor with the grating and the platform. In a failed trial, the mouse crossed the midline divider to the side with the isoluminant gray screen. Each mouse completed up to 50 trials/day in blocks of 10. To advance past training, mice had to achieve two consecutive blocks with 80% accuracy or better. If not, they were excluded. Before testing, the left eye of each mouse was sutured shut (monocular deprivation), and an additional training period was repeated with the same criterion. The mouse had to complete two consecutive blocks with 8/10 correct trials (80%). Typically, this was accomplished in 20-50 trials over 1-2 days.

Testing began with the discrimination between a low frequency grating (0.05 cpd) and an isoluminant screen. After three consecutive correct choices (3/3), the SF was increased by one additional bar on the subsequent trials until the mouse could no longer discriminate between the stimulus and control. Validation of this measure included retesting at the failed SF, requiring either five consecutive correct trials (5/5) or 8/10 correct trials at the failed SF to advance. If unsuccessful (>8/10 correct trials) at the failed SF, retraining at half the SF to correct any possible ‘side bias’ in their performance. The mouse would then resume testing at the SF two steps below the failed SF. Testing ended when mice failed three times at adjacent SFs (i.e, 0.42, 0.47, 0.44). The three failed SFs were averaged and used to define the visual acuity threshold.

Photopic visual acuity was tested with monitor brightness at the choice point was 100 cd/m^2^. Scotopic visual acuity was tested with monitor brightness at the choice point was 0.025 cd/m^2^, which was achieved by lowering the monitor brightness to 25% and using neutral density (ND) filters (ND 0.9, ND 0.9, ND 1.2) after one hour of dark adaptation with room luminance ≤0.01 cd/m^2^. All equipment emitting light was covered, and the experimenter was dark-adapted for 20 minutes and worked under dim red light.

### Electroretinogram (ERG)

Mice were dark-adapted overnight, anesthetized with a solution of ketamine (80mg/kg) and xylazine (16 mg/kg) administered via intraperitoneal injection (IP), and prepared for ffERG recordings under dim red light. Pupils were dilated and accommodation relaxed with the application of eye drops, with 2.5% phenylephrine hydrochloride Ophthalmic Solution (Bausch+Lomb, NDC 82260-102-10) and 0.5% Tropicamide Ophthalmic Solution (Bausch+Lomb NDC 24208-590-64) for 30 seconds. Eyes were then rinsed three times with sterile irrigating solution (Alcon Balanced Salts Solution [BSS]). A contact lens with a gold electrode (LKC Technologies Inc.) was placed on the cornea and position maintained with a hypromellose ophthalmic solution (Vista Gonio Eye Lubricant). Ground and reference needle electrodes were placed in the tail and on the midline of the forehead, respectively. Body temperature was maintained using a feedback-controlled electric heating pad (37°C).

Scotopic responses were measured at two test flash intensities (0.025 and 0.05 cd s/m^2^). After 10 minutes of adaptation to a rod-saturating background (20 cd/m^2^). Photopic responses were measured at two test flash intensities (10 and 100 cd s/m^2^). For each stimulus, a-wave, b-wave amplitudes, and b-wave implicit times were analyzed with a custom MATLAB code. The a-wave was measured from baseline (recorded 10-20 msec before stimulation) to the negative trough. The b-wave was measured from the a-wave trough to the b-wave peak. The flash intensities in the protocol for this experiment, scotopic flash (0.025 cd s/m^2^) and the photopic flash (100 cd s/m^2^), matched the visual water task screen luminance, allowing for direct comparison between behavioral vision acuity and retinal function. ERGs were performed within 2 weeks after concluding measuring acuity with the visual water task.

### Cranial windows

Wide field epi-fluorescent calcium imaging and two-photon calcium imaging were performed though a cranial window as previously described (Brown et al., 2024; Trachtenberg et al., 2002). In brief, mice were administered carprofen (5 mg/kg) and buprenorphine (0.1 mg/kg) for analgesia and anesthetized with isoflurane (4% induction, 1% to 2% maintenance). The scalp was shaved and mice were mounted on a stereotaxic frame with palate bar and their body temperature maintained at 37°C with a heat pad controlled by feedback from a rectal thermometer (TCAT-2LV, Physitemp). The scalp was resected, the connective tissue removed from the skull, and a custom aluminum headbar affixed with C&B metabond (Parkell). A circular region of bone 3 mm in diameter centered over left visual cortex was removed using a high-speed drill (Foredom). Care was taken to not perturb the dura. A sterile 3 mm circular glass coverslip was sealed to the surrounding skull with cyanoacrylate (Pacer Technology) and dental acrylic (Ortho-ject, Lang Dental). The remaining exposed skull likewise sealed with cyanoacrylate and dental acrylic. Mice recovered on a heating pad and returned to standard housing for at least 2 days prior to 2-photon imaging. Implanting the cranial window and imaging were performed within 2 weeks after concluding measuring acuity with the visual water task.

### Wide field epi-fluorescent calcium imaging

After implantation of the cranial window and before 2-photon imaging, the binocular zone of visual cortex was identified with wide field calcium imaging similar to our method for optical imaging of intrinsic signals (Frantz et al., 2016; Kalatsky et al., 2005). In brief, mice were anesthetized with isoflurane (4% induction), provided a low dose of the sedative chlorprothixene (0.5 mg/kg IP; C1761, Sigma) and secured by the aluminum headbar. The eyes were lubricated with a thin layer of ophthalmic ointment (Puralube, Dechra Pharmaceuticals). Body temperature was maintained at 37°C with heating pad regulated by a rectal thermometer (TCAT-2LV, Physitemp). Visual stimulus was provided through custom-written software (MATLAB, Mathworks). A monitor was placed 25 cm directly in front of the animal and subtended +40 to −40 degrees of visual space in the vertical axis. A horizonal white bar (2 degrees high and 20 degrees wide) centered on the zero-degree azimuth drifted from the top to bottom of the monitor with a period of 8 seconds. The stimulus was repeated 60 times. Cortex was illuminated with blue light (475 ± 30 nm) (475/35, Semrock) from a stable light source (Intralux dc-1100, Volpi). Fluorescence was captured utilizing a green filter (HQ620/20) attached to a tandem lens (50 mm lens, Computar) and camera (Manta G-1236B, Allied Vision). The imaging plane was defocused to approximately 200 μm below the pia. Images were captured at 10 Hz as images of 1,024 × 1,024 pixels and 12-bit depth. Images were binned spatially 4 × 4 before the magnitude of the response at the stimulus frequency (0.125 Hz) was measured by Fourier analysis.

### Visual stimuli and two-photon calcium imaging

Visual stimulus presentation and image acquisition were both performed according to our published methods which were modified from published studies (Brown et al., 2024; Brown and McGee, 2023; Jimenez et al., 2018; Tan et al., 2020). In brief, a battery of static sinusoidal gratings was generated in real time with custom software (Processing, MATLAB). Stimulus presentation was synchronized to the imaging data by time stamping the presentation of each visual stimulus to the image acquisition frame number a transistor–transistor logic (TTL) pulse generated with an Arduino at each stimulus transition. Orientation was sampled at equal intervals of 15 degrees from 0 to 150 degrees (12 orientations). SF was sampled in 10 steps on a logarithmic scale at half-octaves from 0.028 to 1.02 cpd. An iso-luminant grey screen was included (blank) was provided as a ninth step in the SF sampling as a control. Spatial phase was equally sampled at 45-degree intervals from 0 to 315 degrees for each combination of orientation and SF. Gratings with random combinations of orientation, SF, and spatial phase were presented at a rate of 4 Hz on a monitor with a refresh rate of 60Hz. Imaging sessions were 20 minutes (4800 gratings presented in total). Consequently, each combination of orientation and SF was presented 33 times on average (range 17 to 56). The monitor was centered on the zero azimuth and elevation 35 cm away from the mouse and subtended 45 (vertical) by 80 degrees (horizontal) of visual space.

Imaging was performed with a resonant scanning 2-photon microscope controlled by Scanbox image acquisition and analysis software (Neurolabware). The objective lens was fixed at vertical for all experiments. Fluorescence excitation was provided by a tunable wavelength infrared laser (Ultra II, Coherent) at 920 nm. Images were collected through a 16× water-immersion objected (Nikon, 0.8 NA). Images (512 × 796 pixels, 520 × 740 μm) were captured at 15.5 Hz at depths between 150 and 400 μm. Eye movements and changes in pupil size were recorded using a Dalsa Genie M1280 camera (Teledyne Dalsa) fitted with 50 mm 1.8 lens (Computar) and 800 nm long-pass filter (Edmunds Optics). Imaging was performed on alert mice positioned on a spherical treadmill by the aluminum head bar affixed to the skull. The visual stimulus was presented to the contralateral eye by covering the fellow eye with a small custom occluder. Neutral density filters were positioned in front of the monitor presenting visual stimuli for imaging under of neuronal activity under scotopic conditions. Mice were dark-adapted for 30 minutes or longer prior to imaging. Mice were imaged within two weeks of measuring visual acuity with the visual water task.

### Image Processing

Image processing was performed as described previously (Brown and McGee, 2023; Tan et al., 2020). Imaging series for each eye were motion corrected with the SbxAlign tool. Regions of interest (ROIs) corresponding to excitatory neurons were selected manually with the SbxSegment tool following computation of pixel-wise correlation of fluorescence changes over time from 350 evenly spaced frames (~1%). ROIs for each experiment were determined by correlated pixels the size similar to that of a neuronal soma. The fluorescence signal for each ROI and the surrounding neuropil were extracted from this segmentation map.

### Image Analysis

Image analysis was performed as described previously with minor modifications (Brown and McGee, 2023). The fluorescence signal for each neuron was extracted by computing the mean of the calcium fluorescence within each ROI and subtracting the median fluorescence from the surrounding perimeter of neuropil (Ringach et al., 2016; Tan et al., 2020). An inferred spike rate (ISR) was estimated from adjusted fluorescence signal with the Vanilla algorithm (Berens et al., 2018). A reverse correlation of the ISR to stimulus onset was used to calculate the preferred stimuli (Brown and McGee, 2023; Jimenez et al., 2018; Ringach et al., 2016; Tan et al., 2020). Neurons that satisfied 3 criteria were categorized as visually responsive: (1) the ISR was highest with the optimal delay of 4 to 9 frames following stimulus onset. This delay was determined empirically for this transgenic GCaMP6s mouse (Brown and McGee, 2023; Tan et al., 2020); (2) the SNR was greater than 1.3. The signal is the mean of the spiking standard deviation at the optical delay between 4 and 9 frames after stimulus onset and the noise this value at frames −2 to 0 before the stimulus onset or 15 to 18 after it (Jimenez et al., 2018; Tan et al., 2020). (3) and neuron responded to at least 13% of the presentations of the preferred stimulus. Visual responsiveness for every neuron was determined independently for each eye. The visual stimulus capturing the preferred orientation and SF was the determined from the matrix of all orientations and SFs presented as the combination with highest average ISR.

The preferred orientation for each neuron was calculated as:

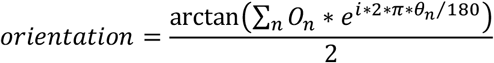

O_n_ is a 1 × 12 array of the mean z-scores associated with the calculation of the ISR at orientations Q_n_ (0 to 150 degrees, spaced every 15 degrees). Orientation calculated with this formula is in radians and was converted to degrees. The tuning width was the full width at half-maximum of the preferred orientation.

The preferred SF for each neuron was calculated as:

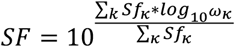

Sf_k_ is a 1 × 10 array of the mean z-scores at SFs w_k_ (10 equal steps on a logarithmic scale from 0.028 to 1.02. cpd). Tails of the distribution were clipped at 25% of the peak response. The tuning width was the full width at half-maximum of the preferred SF in octaves. The percent visually responsive neurons with significant responses at each SF was determined by comparing the distribution of ISR values at each SF versus the stimulus blank with a KW-test with Dunn’s correction for 10 comparisons. Neurons with P < 0.01 for a given SF were considered significant responses at that SF (Salinas et al., 2017).

### Retinal Dissection

The mouse was euthanized with a solution of ketamine (80mg/kg) and xylazine (16 mg/kg) administered via intraperitoneal injection (IP). After a toe pinch evoked no reaction, cervical dislocation was performed as a secondary form of euthanasia. The superior portion of the right eye was marked using a pen (Securline Surgical Skin Marker), and the eye was enucleated. A hole punched with an #11 scalpel and a circumferential cut separated the cornea and lens from the eyecup. The retina was removed after cutting the optic nerve and separating it from the posterior eye cup. One deep cut was made at the mark on the superior retina, and two more minor cuts were made at the nasal and temporal sides of the inferior retina. This allowed the retina to be flattened on a glass slide where it was fixed with 4% paraformaldehyde in phosphate-buffered saline (PBS) solution, pH 7.4, for 20 minutes, then washed with PBS. An ascending sucrose series was used to cryoprotect the retinal whole mounts (5%, 10%, 15%, one hour each; 20%(sucrose/PBS) overnight). The whole mount retina was placed in 2:1 Sakura Tissue-Tek O.C.T. Compound (Product Code 4583)/20% sucrose for an hour, embedded in the same solution, and frozen in a 2-methyl butane liquid nitrogen-cooled bath. Retinas were sectioned at 14 µm using a Leica 1850 cryostat, mounted on Superfrost Plus glass slides (Fisher Scientific 12-550-15), dried for 60 minutes on a heating plate (37°C), and then stored at −80°C.

### Histochemistry

Slides were thawed on a heating plate (37°C) for 50-60 minutes and sections outlined on the slide with a hydrophobic barrier (PAP pen; Vector Laboratories H-4000). Each slide was washed with ~200uL of 1x PBS (Invitrogen AM9625) for 5 minutes and sections permeabilized with 0.5% TritonX-100 (MP Biomedicals ICN807423) in 1x PBS. A blocking solution of 1x PBS with 5% NDS (Normal Donkey Serum, Millipore Sigma S30-100ML) was applied, and slides were incubated for 1 hour at room temperature followed a solution containing DAPI (4’,6-Diamidino-2-Phenylindole, Invitrogen, D1306; 1:2000) to stain somas.

### Outer Nuclear Layer Measurements

Images of the retinal sections were taken on a confocal laser scanning microscope (Olympus FV4000). A stitched image of the entire section was acquired, and high-power images were acquired at 100 µm intervals. Across the section, at each area, a Z stack of 1 µm images (14 µm total) was acquired using a 40x objective with a 1.5x zoom. Olympus cellSens Dimensions software measured the retinal layers at each area from a projection consisting of the middle five images of the Z stack, ~5µm depth, the diameter of photoreceptor somas. Six evenly spaced measurements of the ONL were made across each image (superior to inferior). All measurements for a single retina (6 images x 6 measurements = 36 measurements) were averaged to define the average ONL thickness.

### Statistics

No statistical methods were used to predetermine sample size. All statistical analyses were done using Prism 8 software (GraphPad Software). Multiple comparisons were tested with Analysis of Variance (ANOVA) tests and paired tests with Mixed-effects analysis for data not divergent from a parametric distribution and the non-parametric Kruskal-wallis test for data that did not fit a parametric distribution.

## Acknowledgements

This work was supported by a grant from the National Eye Institute (EY035885 to AWM and MAM) and a Jewish Heritage Fund for Excellence Research Enhancement Grant (MAM).

## Author contributions

CAA, TCB, MAM, and AWM conceived and designed the study. CAA and TCB performed experiments. CAA, TCB, MAM, and AWM analyzed the data. CAA, TCB, MAM, and AWM wrote the manuscript.

## Competing interests

The authors have declared that no competing interests exist.

## Notes

### Competing Interest Statement

The authors have declared no competing interest.

