## Supplemental Figures for "Enhancing experience-dependent plasticity accelerates vision loss in a murine model of retinitis pigmentosa"

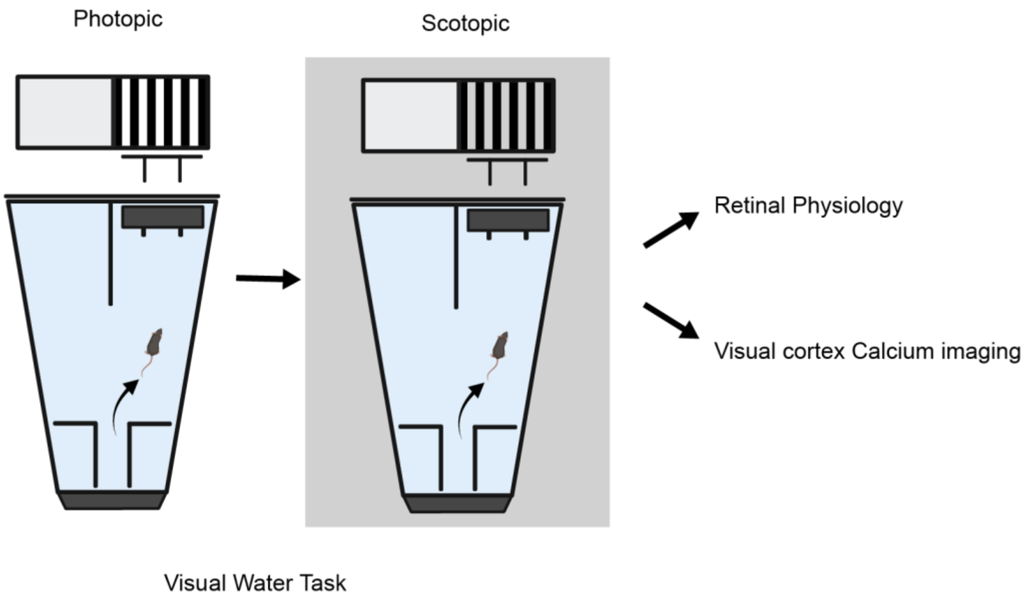


**Supplemental Figure 1. Overview of the experimental paradigm.**
Mice are first tested to measure acuity with the visual water task under scotopic conditions for the right eye (monocular vision) and then retested under scotopic conditions. Thereafter, mice that are transgenic and express GCaMP6s in forebrain excitatory neurons are directed to *in vivo* calcium imaging at neuronal resolution and non-transgenic mice are routed to measurements of retinal physiology.

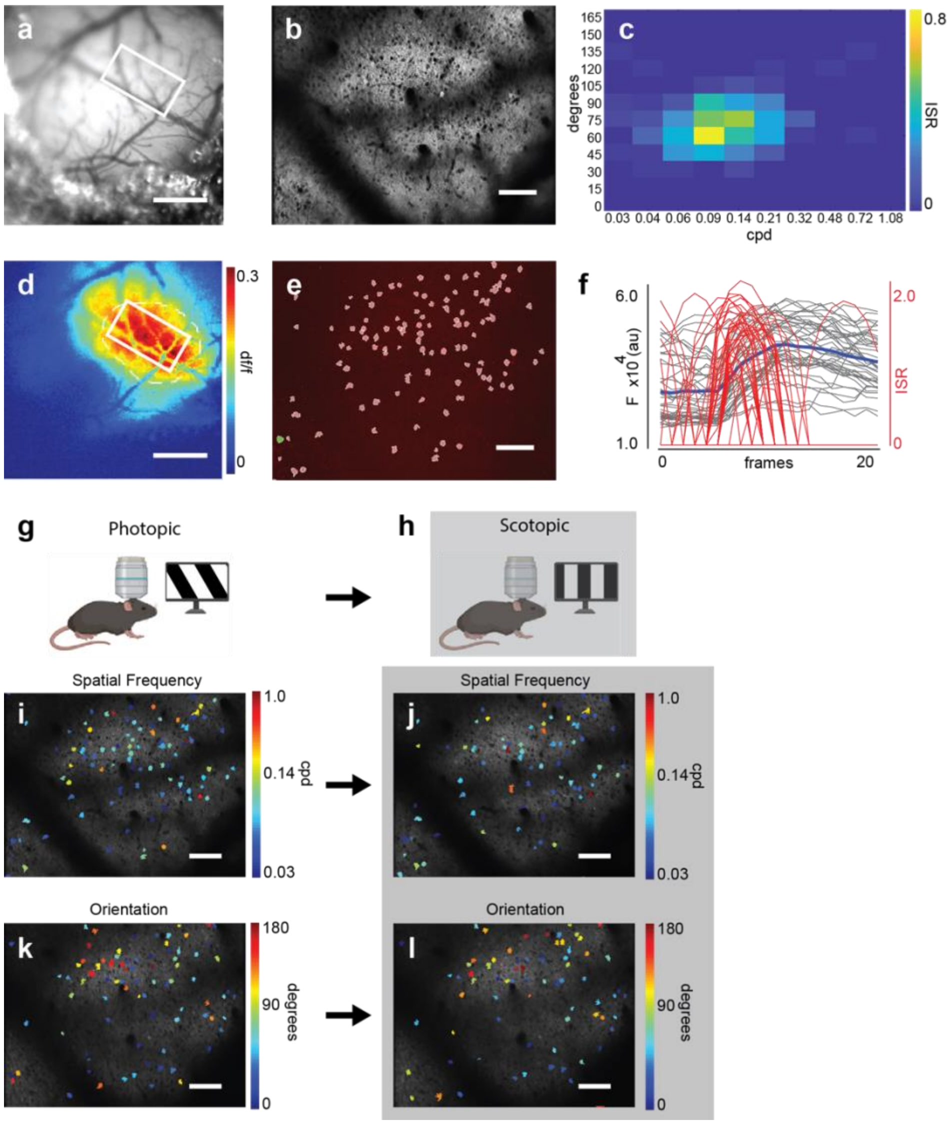


**Supplemental Figure 2. Two-photon calcium imaging of visual cortex under photopic and scotopic conditions.** **(a)** Example cranial window. A 3 mm glass coverslip is placed over visual cortex. The white rectangle indicates the imaging field. Scale bar = 1mm. **(b)** Example imaging field of visual cortex. Scale bar = 100μm. **(c)** Example heat map of inferred spike rate (ISR) for representative neuron in imaging field. Stimulus orientation is on the y-axis and stimulus spatial frequency is on the x-axis (cpd, cycles per degree). **(d)** Wide field calcium of visual cortex. Red colors indicate the region of greatest response magnitude. Scale bar = 1mm. **(e)** Segmentation map of regions of interest (ROIs) from imaging field. **(f)** Fluorescent traces (grey lines) superimposed for the 20 frames (1.25 seconds) following the onset of the 39 presentations of the preferred visual stimulus for an example neuron. Red lines represent the positions of inferred spikes. The blue line indicates the average fluorescence across all frames and presentations. **(g,h)** Cartoon of 2-photon imaging set up during photopic (panel g) and scotopic conditions (panel h). Awake mouse head fixed and freely running on styrofoam ball is imaged while 4hz stimulus is presented. For imaging under dark adapted conditions, room lights are turned off and neutral density filters are placed over monitor presenting the visual stimulus. **(i,j)** Preferred spatial frequency tuning for neurons in an imaging field under light-adapted (i) and dark-adapted (j) conditions. Scale bar = 100μm. **(k,l)** Preferred orientation tuning for neurons in an imaging field under light-adapted (k) and dark-adapted (l) conditions. Scale bar = 100μm.

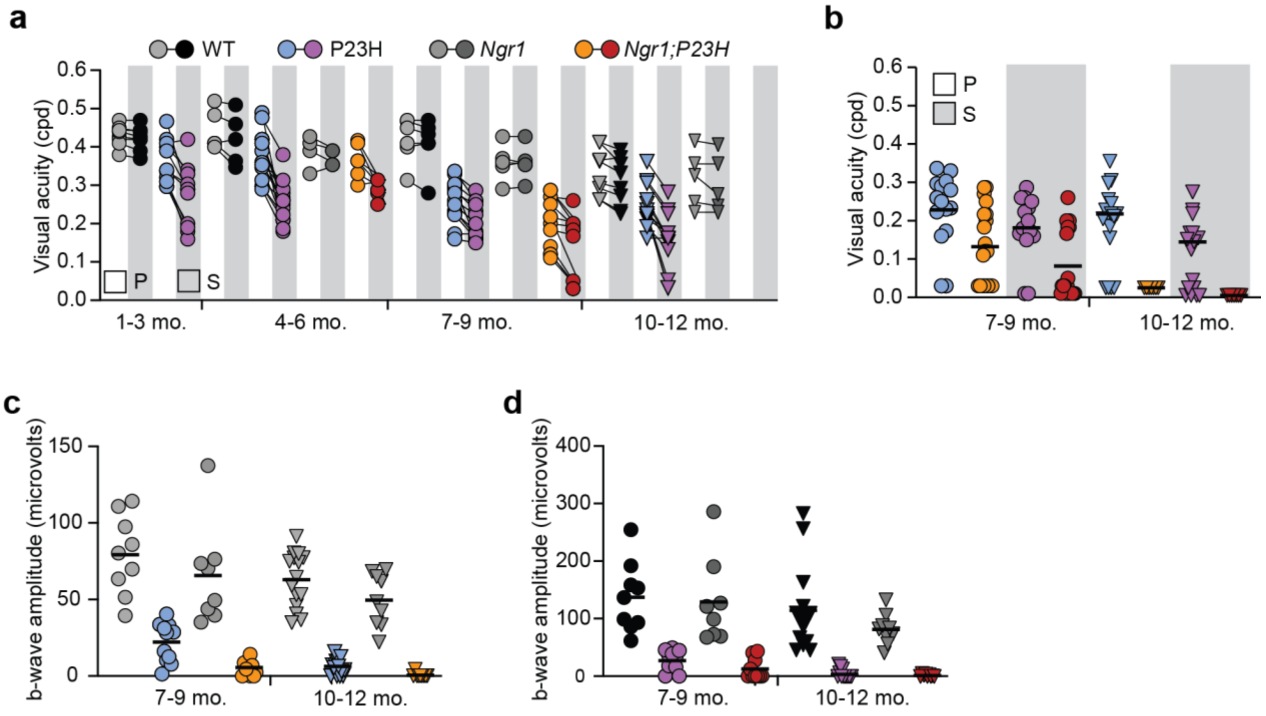


**Supplemental Figure 3. Comparison of acuity and ERG b-wave amplitudes under photopic and scotopic conditions for WT mice, *Rho P23H/+* mice, *Ngr1-/i* mice; and *Ngr1-/-; Rho P23H/+* mice. (a)** Acuity measured under photopic (P) and scotopic conditions (S, grey rectangles) for wild-type (WT), *Rho P23H/+* (P23H), *Ngr1-/-* (*Ngr1*), and *Ngr1-/-; Rho P23H/+* (*Ngr1;P23H*) mice. Circles represent mice at 7-9 mo. and triangles at 10-12 mo.. Lines connect measurements under each luminance condition. Color scheme for genotype and luminance conditions are indicated above the plot. Ages tested are indicated at bottom in months (mo.) of age. cpd, cycles per degree. Data presented is reorganized from Fig. 1a and Fig. 4a. **(b)** Column plot of acuity for 7-9 mo. and 10-12 mo. Color and symbol scheme for genotype, luminance conditions, and age are the same as panel a. **(c,d)** b-wave amplitudes under photopic conditions (panel c) and scotopic conditions (panel d) for the genotypes presented in panel a. Data presented is reorganized from Fig. 2c,d and Fig. 4c,d. Color and symbol scheme for genotype, luminance conditions, and age are the same as panel a.

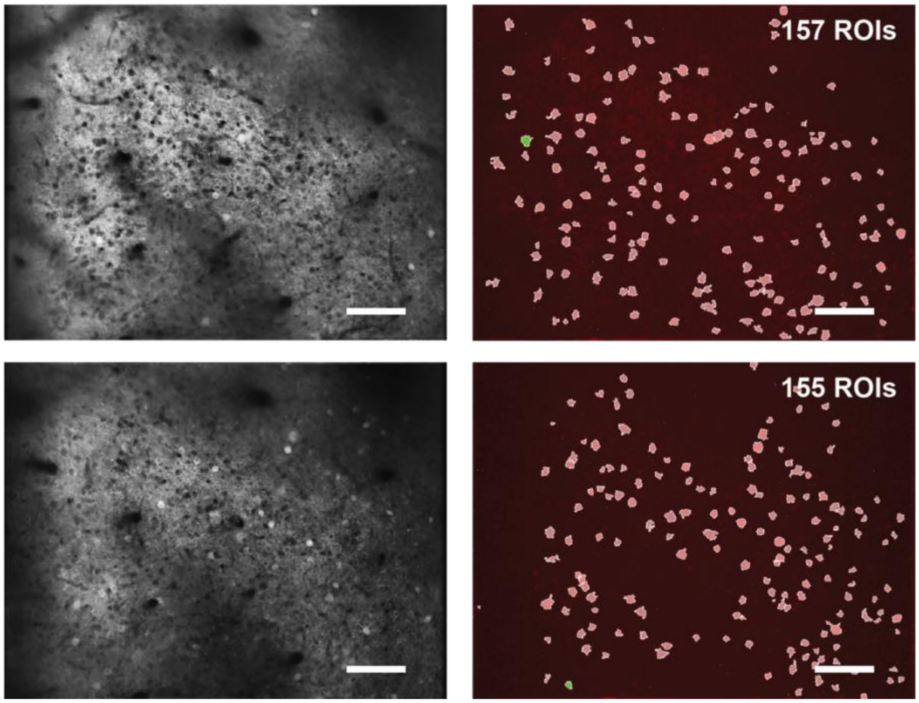


**Supplemental Figure 4. Example imaging fields from functionally blind *Ngr1 -/-; Rho P23H/*+ mice.**
Two imaging fields from visual cortex of an *Ngr1 -/-; Rho P23H/*+ mouse that did not meet threshold criteria for testing on the visual water task. Reference images are on the left and segmented regions of interest are on the right. Scale bar = 100μm. The low yield of visually responsive neurons was not a consequence of a poor-quality cranial window or lack of identifiable neurons.
